# Mechanosensitive clathrin platforms anchor desmin intermediate filaments in skeletal muscle

**DOI:** 10.1101/321885

**Authors:** Agathe Franck, Jeanne Lainé, Gilles Moulay, Michaël Trichet, Christel Gentil, Anaïs Fongy, Anne Bigot, Sofia Benkhelifa-Ziyyat, Emmanuelle Lacène, Mai Thao Bui, Guy Brochier, Pascale Guicheney, Sabrina Sacconi, Vincent Mouly, Norma Romero, Catherine Coirault, Marc Bitoun, Stéphane Vassilopoulos

## Abstract

Large flat clathrin plaques are stable features of the plasma membrane associated with sites of strong adhesion suggesting that they could also play a role in force transduction. Here, we analyzed how clathrin plaques interact with the cytoskeleton and how they respond to mechanical cues in skeletal muscle myotubes. We show that branched actin networks surrounding clathrin plaques are directly regulated by dynamin 2, anchor intermediate filaments and sequester YAP at the plasma membrane. Dynamin 2, clathrin and desmin intermediate filaments are all required for basal YAP nucleocytoplasmic distribution and efficient nuclear translocation in response to mechanical stimuli. Dynamin 2 mutations that are responsible for centronuclear myopathy in humans disorganize the desmin network and deregulate YAP signaling both *in vitro* and *in vivo*. Thus, clathrin plaques and associated dynamin 2 are defined here as a new sensor conveying mechanical cues and integrate cell signaling with cytoskeletal regulation.

## Introduction

For vesicle formation, clathrin triskelia composed of trimerized heavy chains (CHCs) with bound light chains (CLCs), are recruited by clathrin adaptors that trigger clathrin-coated vesicle (CCV) budding (Brodsky, 2012; Robinson, 2015). The adaptor proteins are required for targeting clathrin to specific intracellular compartments and, among these, adaptor protein 2 (AP2) recruits clathrin to the plasma membrane (PM). In several cell types, and most notably in skeletal muscle myotubes, flat clathrin plaques cover large portions of the PM (Grove et al., 2014; Heuser, 1980; Lampe et al., 2016; Maupin and Pollard, 1983; Saffarian et al., 2009; Taylor et al., 2011). Although flat clathrin lattices are thought to be a structural intermediate which will bud to form a canonical coated pit (Avinoam et al., 2015; Heuser, 1980), the role of these extensive flat clathrin plaques remains elusive. They are invariantly associated with the actin cytoskeleton (Heuser, 1980; Leyton-Puig et al., 2017; Saffarian et al., 2009; Vassilopoulos et al., 2014), they mediate adherence to the extracellular substrate through integrins such as integrin β5 (De Deyne et al., 1998; Lampe et al., 2016; Vassilopoulos et al., 2014) and are thought to serve as hot spots for endocytosis since budding pits are frequently observed at their periphery (Lampe et al., 2016; Taylor et al., 2011). Dynamin 2 (DNM2), a large GTPase which acts as a mechanochemical scaffolding molecule that releases nascent vesicles from the PM or intracellular compartments (van Dam and Stoorvogel, 2002; Kaksonen and Roux, 2018), is a *bona fide* clathrin plaque component (Grove et al., 2014). In addition, several studies have demonstrated that DNM2 regulates actin cytoskeleton networks and have proposed functions for dynamin in actin polymerization that are distinct from coated vesicle formation (Bonazzi et al., 2012; González-Jamett et al., 2017; Gu et al., 2010; Orth and McNiven, 2003; Saffarian et al., 2009; Schafer et al., 2002). Mutations in the *DNM2* gene cause autosomal dominant centronuclear myopathy (CNM) (Bitoun et al., 2005), a slowly progressive congenital myopathy which is characterized by muscle weakness, hypotonia and the presence of centrally located nuclei in a large number of muscle fibers in the absence of regenerative processes.

In skeletal muscle, CHC and DNM2, are localized at specific PM sites called costameres. CHC depletion *in vivo* leads to drastic detachment of peripheral contractile apparatus from the plasma membrane and decreased force (Vassilopoulos et al., 2014). Costameres correspond to lateral contacts between the PM and the contractile apparatus (Pardo et al., 1983b, 1983a; Shear and Bloch, 1985) and play a pivotal role integrating adhesion to the propagation of forces (Danowski et al., 1992). They are composed of large membrane protein complexes, i.e. the focal adhesion complex and the dystrophin-glycoprotein complex that are both linked to the contractile apparatus by actin and desmin intermediate filaments (IFs) (Ervasti, 2003). Desmin is a muscle-specific, type III intermediate filament protein, well documented for its role in cell architecture and force transmission (Capetanaki et al., 2015; Herrmann et al., 2007), which forms a stress-transmitting network involved in signalling, mechanotransduction and gene expression (Palmisano et al., 2015). Costameres are bidirectional mechanochemical signal transduction sites where contractile forces generated within the fiber are directly transmitted to the surrounding extracellular matrix (ECM) and where longitudinal displacement of the ECM is transmitted to the contractile machinery (Samarel, 2005). Although it is established that mechanical cues sensed by costameres are transduced into biochemical signals leading to sarcomere assembly and gene expression regulation, little is known on specific mechanisms involved. Among them, two recently identified key mechanotransduction players, Yes-associated protein (YAP) (Sudol, 1994) and Transcriptional co-activator with PDZ-binding motif (TAZ) (Dupont et al., 2011; Kanai et al., 2000), are transcriptional cofactors that can shuttle between the cytoplasm and the nucleus in response to mechanical cues from the ECM (Aragona et al., 2013; Cui et al., 2015; Halder et al., 2012). Although the precise nature of the compartments that YAP and TAZ associate with in the cytoplasm is unknown, their activity depends on cytoplasmic F-actin structures (Aragona et al., 2013).

Here, we set out to investigate the mechanism by which adhesive clathrin plaques could organize a compartment capable of responding to mechanical cues. We report clathrin plaques and the associated branched actin filaments are required to anchor desmin IFs at costameres and form signaling platforms for YAP. Alteration of this clathrin-based mechanotransducing platform by *DNM2* mutations which are responsible for CNM highlights a novel pathophysiological mechanism of this congenital myopathy.

## Results

### Clathrin plaques act as platforms for cytoskeletal organization

We analyzed clathrin plaques from extensively differentiated primary mouse myotubes. Clathrin-positive fluorescent clusters periodically aligned along the PM following organized striated peripheral α-actinin-positive structures (Fig. 1A). We developed a metal-replica electron microscopy (EM) approach aimed at visualizing these structures *en face* from highly differentiated unroofed myotubes. Platinum replicas obtained from both primary mouse (Fig. 1B-D) and human (Supplementary Fig. 4A-C) myotubes presented regularly spaced clathrin plaques interacting with cortical cytoskeletal filaments. Three-dimensional interaction between cytoskeletal components surrounding clathrin plaques was defined by a combination of platinum replica EM and electron tomography by collecting images at the tilt angles up to ±25° with 5° increments relative to the plane of the sample (Supplemental Movie 1). The small clusters of branched actin surrounding clathrin plaques formed a clamp for thicker rope-like filaments weaving through the actin network (Supplementary Fig. 1A, B) and displayed ArpC2 immunogold labeling (Supplementary Fig. 1C, D). Direct visualization and measurements of filament diameter at the ultrastructural level on 3D anaglyphs allowed us to unambiguously discriminate between straight and thin 10 nm on average actin filaments versus thicker 15 nm filaments (including metal coating). Using immunogold labelling combined to metal-replica EM we showed that this three-dimensional network directly connecting actin stress fibers and branched actin "nests" surrounding flat clathrin plaques is composed of desmin IFs (Fig. 1E and Supplementary Fig. 1E, F). We tested the possibility that clathrin plaques were required for organization of the cortical desmin network. CHC depletion in myotubes induced a strong aggregation of desmin in the cytoplasm (Fig. 1F, G). In order to prove that the phenotype was due to clathrin recruited specifically at the PM, we depleted the α-subunit of the AP2 adaptor, which phenocopied CHC depletion and induced the formation of desmin aggregates in the cytoplasm (Fig. 1G). Importantly, depleting AP1 (γ-subunit) or AP3 (δ-subunit), involved in the recruitment of clathrin to the Golgi apparatus and endosomal systems respectively, had no effect on desmin distribution (Fig. 1G). Thin-section EM confirmed that the aggregates induced by CHC or AP2 depletion were composed of IF tangles (Fig. 1H-O). We tested whether desmin was required for clathrin plaque formation by culturing primary myotubes from desmin knock-out mice. These myotubes displayed regularly spaced plaques that were indiscernible from those observed in WT mouse myotubes (Supplementary Fig. 2). Thus, clathrin plaques and branched actin are necessary for organizing desmin IFs while clathrin plaques can form in the absence of desmin. Altogether, these data demonstrate the importance of clathrin plaques and the surrounding branched actin network for proper organization of the intermediate filament web and suggest that their initial formation is a prerequisite for desmin scaffolding.

**Fig. 1.**
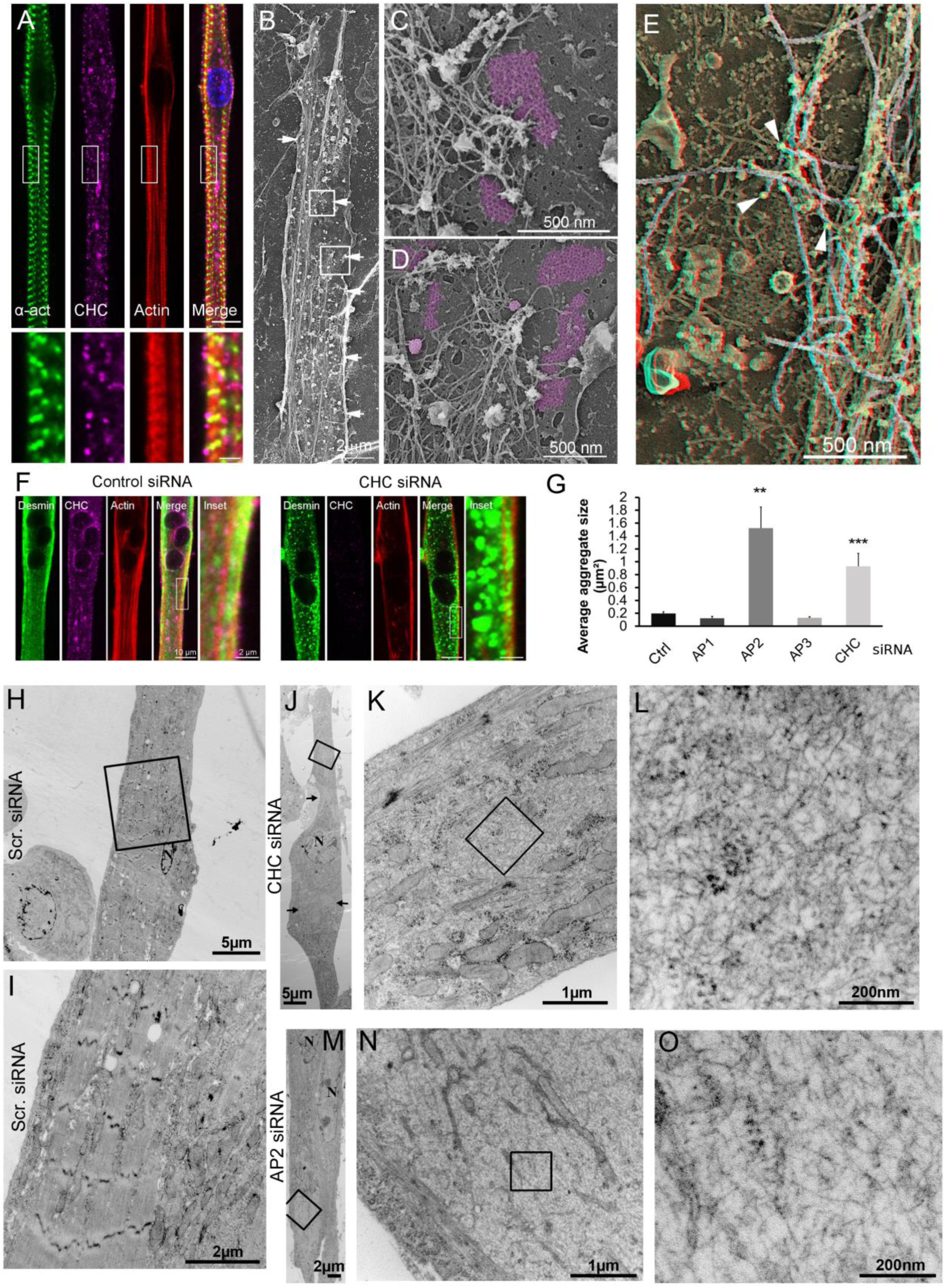
Clathrin-coated plaques are required for intermediate filament organization. **(A)** Immunofluorescent staining of α-actinin 2 (green), CHC (magenta), and actin (red) in extensively differentiated mouse primary myotubes. Bars are 10 µm and 2 µm for insets. **(B)** Survey view of unroofed primary mouse myotube differentiated for 15 days. **(C) (D)** Higher magnification views corresponding to the boxed regions in b. **(E)** Unroofed control primary myotubes were labelled with desmin antibodies (pseudocolored yellow, arrowheads). Intermediate filaments are highlighted in grey. Use glasses for 3D viewing of anaglyph (left eye = red). **(F)** Immunofluorescent staining of desmin (green), CHC (magenta) and actin (red) in mouse primary myotubes treated with control (left) siRNA or siRNA against CHC (right). **(G)** Average desmin aggregate size in myotubes treated with control siRNA or siRNA against CHC, AP1, AP2 or AP3 (*n* = 30–50 myotubes. Data presented as mean ± SEM; **P < 0.01, ***P < 0.001, ****P < 0.0001). **(H-O)** Thin-section EM of primary myotubes treated with control (H-I), CHC (J-L) or AP2 (M-O) siRNA. L and O are higher magnification views of desmin tangles from (K) and (N) respectively.

### Branched actin surrounding clathrin plaques is regulated by N-WASP and DNM2 and organizes desmin intermediate filaments

Neuronal Wiskott–Aldrich Syndrome protein (N-WASP) is directly involved in the generation of Arp2/3 branched actin filaments during endocytosis and a known DNM2 indirect partner (Takenawa and Suetsugu, 2007). N-WASP depletion induced desmin aggregates similar to those produced by CHC depletion (Fig. 2A, C and Supplementary Fig. 3, B-D). DNM2-depleted myotubes also consistently displayed desmin aggregates in the cytoplasm (Fig. 2B, D, H) as well as increased clathrin patches on the myotube's lateral membrane (Fig. 2B, E). Given DNM2 function in actin remodeling, we reasoned that it could participate to organize branched actin filaments including those surrounding clathrin plaques which anchor intermediate filaments. We observed an interaction between DNM2 and N-WASP and some desmin upon DNM2 immunoprecipitation (Fig. 2F). At the EM level, DNM2-depleted myotubes consistently displayed desmin aggregates in the cytoplasm (Fig. 2 G, H) and both N-WASP and DNM2 depletion produced characteristic actin accumulations associated with additional proteinaceous material (Fig. 2 I-K and Supplementary Fig. 3B-H), confirming their requirement for regulating branched F-actin dynamics around clathrin plaques. Overall, we showed the central role played by N-WASP and DNM2 in this unappreciated structural facet of clathrin plaques as organizers of the cortical desmin intermediate filament cytoskeleton.

**Fig. 2.**
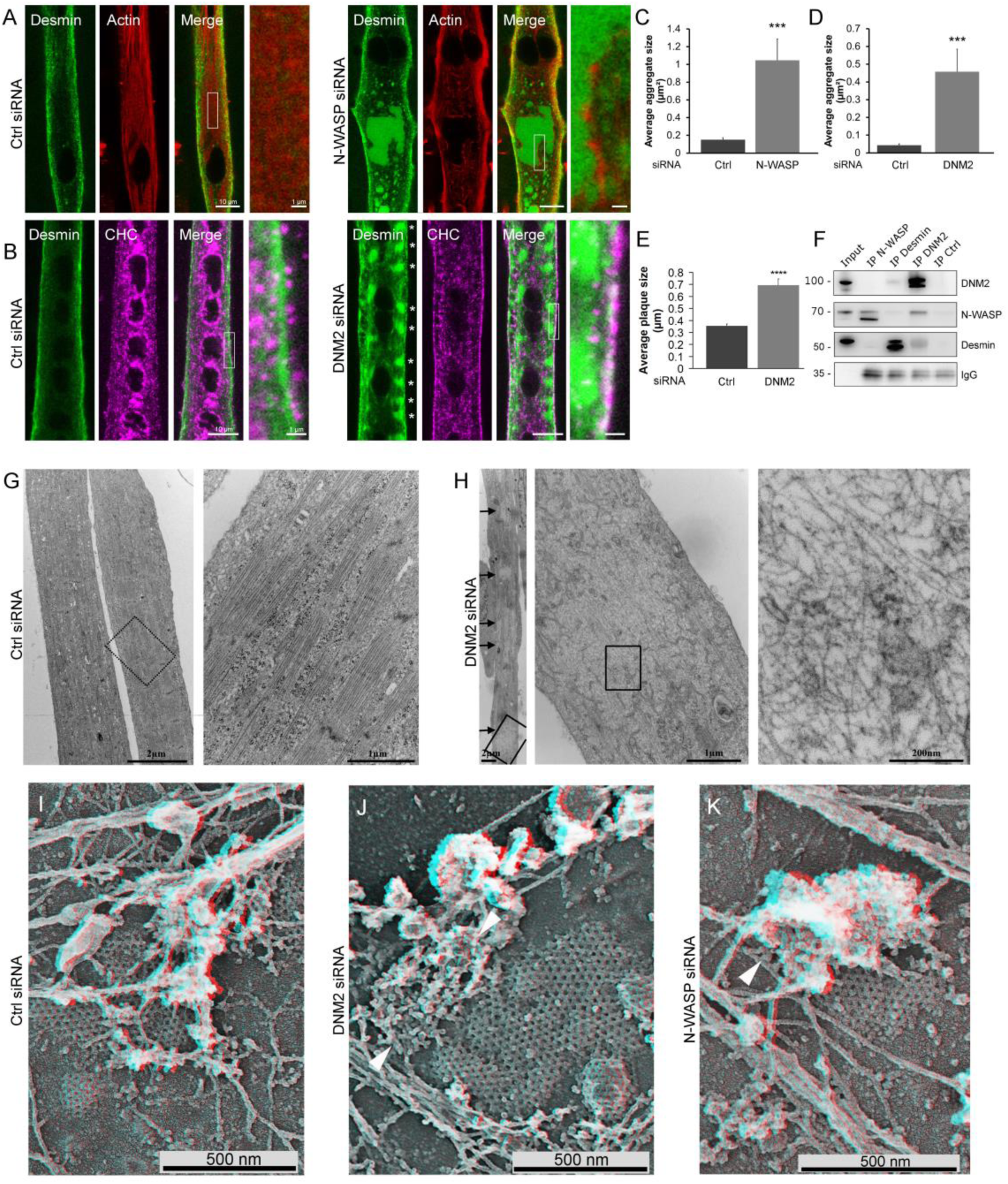
N-WASP and DNM2 are indispensable for desmin and actin organization around clathrin plaques. **(A)** Immunofluorescent staining of desmin (green) and actin (red) in mouse primary myotubes treated with control or N-WASP siRNA. **(B)** Immunofluorescent staining of desmin (green) and CHC (magenta) in control or DNM2-depleted mouse primary myotubes. Asterisks denote desmin aggregates aligned along the PM. **(C) (D)** Desmin aggregate fluorescence in myotubes treated with control, N-WASP (C) or DNM2 (D) siRNA (*n* = 20 myotubes). **(E)** Average CCS size in myotubes treated with control or siRNA against DNM2 (*n* = 20 myotubes). **(F)** Immunoblot of proteins associated with N-WASP, desmin, DNM2, or control immunoprecipitates from mouse primary myotubes lysates. **(G) (H)** Thin-section EM of extensively differentiated control (G) or DNM2-depleted (H) myotubes. Arrows denote clearer IF bundles. **(I-K)** High magnification view of unroofed primary mouse myotubes treated with control (I), DNM2 (J) or N-WASP (K) siRNA. Abnormal actin structures are indicated using arrowheads. Use glasses for 3D viewing of anaglyphs (left eye = red). Data are presented as mean ± SEM; *P < 0.05, **P < 0.01, ***P < 0.001, ****P < 0.0001.

### Clathrin plaques are mechanosensitive platforms and scaffold YAP via surrounding branched actin structures

Given desmin IFs network function in mechanotransduction, we assessed if clathrin-coated structures (CCS) could respond to mechanical cues. We subjected differentiated myotubes grown on a flexible polydimethylsiloxane (PDMS) substrate to stretching/relaxation cycles. Although metal-replica EM is not compatible with myotubes grown on PDMS, we used a spinning-disc confocal microscope equipped with a super-resolution module with a resolution of 150 nm that is sufficient to distinguish clathrin-coated pits from larger plaques (Fig. 3A). By measuring the size of fluorescently labeled AP2 patches at the surface of myotubes, we observed a significant decrease in plaque size with stretching (Fig. 3B). To test if this size reduction was due to increased clathrin-mediated endocytosis rates, we performed a transferrin uptake assay on cyclically stretched myotubes. Transferrin internalization was significantly increased in cells subjected to a cyclic mechanical strain, reflecting higher rates of clathrin-mediated endocytosis (Fig. 3C). Thus, increased clathrin-mediated endocytosis added to plaque size reduction after stretching suggest that clathrin plaques are sensitive to mechanical cues and that prolonged mechanical stimulation could induce plaque disassembly. Mechanical cues such as cyclic stretching cause YAP translocation into the nucleus in undifferentiated myoblasts (Cui et al., 2015; Fischer et al., 2016). Due to their adhesive nature, we reasoned that clathrin plaques could serve as YAP platforms for signaling which could respond to cyclic stretching by increased endocytosis. We used immunogold labeling on metal replicas from unroofed myotubes to specify YAP distribution at the PM. YAP was enriched around plaques as it associated unambiguously with branched actin filaments located around clathrin-coated plaques and clathrin-coated pits in both mouse and human myotubes (Fig. 3, D-H). Half of all the gold beads were directly associated with clathrin lattices or actin structures 100 nm away from clathrin lattices (Fig. 3I). Altogether, these data suggest that clathrin plaques could respond to mechanical cues by their ability to organize a three-dimensional cytoskeletal network around them which acts as a reservoir for YAP signaling.

**Fig. 3.**
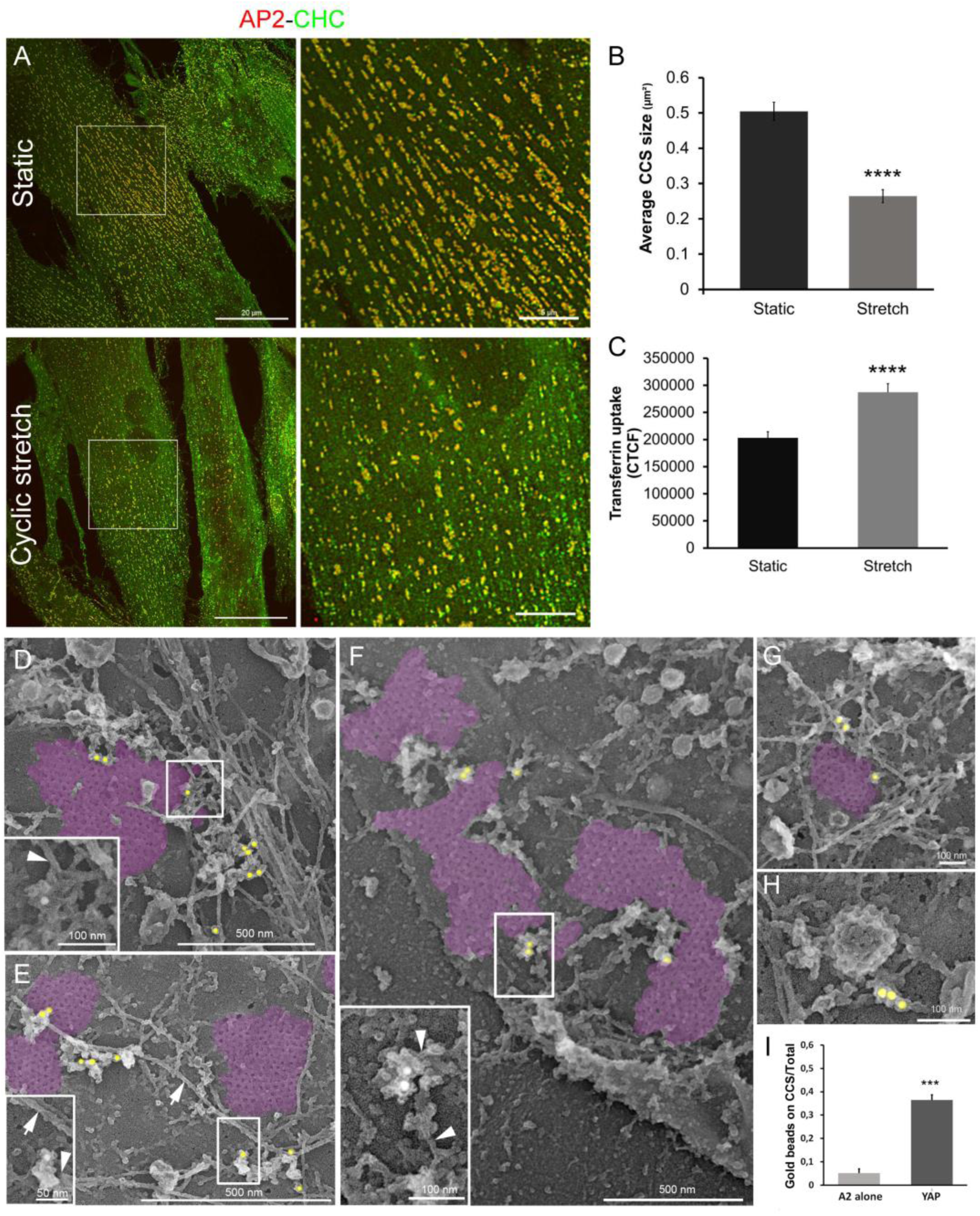
Clathrin platforms are sensitive to cyclic stretching and directly scaffold YAP. **(A)** Immunofluorescent staining of AP2 (red) and CHC (green) in immortalized human myotubes either stretched or left static. **(B)** Average AP2 patch size in human control myotubes either stretched or left static (*n* = 20 myotubes). **(C)** Internalized transferrin in human control myotubes either stretched or left static (*n* = 90–100 myotubes). **(D-H)** High magnification views of clathrin plaques from primary mouse myotubes (D-E) or human myotubes (F) labelled with YAP antibodies (pseudocolored yellow). Clathrin lattices are highlighted in purple. YAP associated to actin cytoskeleton are shown using arrowheads. **(G-H)** High magnification views of clathrin coated pits from primary mouse (D, E, G and H) or human (F) myotubes labelled with YAP antibodies (pseudocolored yellow). **(I)** Number of gold beads corresponding to YAP labeling on clathrin-coated structures or surrounding actin cytoskeleton (<100 nm or closer from the clathrin lattice) compared to a staining performed using only secondary antibodies conjugated to 15-nm colloidal gold particles (*n* = 49 to 104 images). Data presented as mean ± SEM; *P < 0.05, **P < 0.01, ***P < 0.001, ****P < 0.0001.

### CHC, desmin and DNM2 are required for YAP cytoplasmic sequestration and translocation

To test YAP-mediated cellular responses to mechanical cues in differentiated cells, we subjected myotubes grown on a flexible substrate to cyclic stretching under conditions that would induce translocation of YAP (Cui et al., 2015). At the basal state, YAP was mostly cytoplasmic and nuclear staining of YAP increased significantly upon cyclic stretching (Fig. 4A). To test the contribution of clathrin plaques on YAP translocation, we depleted CHC in differentiated myotubes. At the basal state, YAP was strongly accumulated in nuclei from CHC-depleted myotubes (Fig. 4A). Consequently, no further nuclear YAP increase was observed upon cyclic stretching. Primary myotubes from desmin knock-out mice presented a similar accumulation of YAP in myonuclei at the basal state and no further translocation upon stretching (Fig. 4B). We next tested the effect of DNM2 on YAP translocation in myotubes. DNM2-depletion phenocopied CHC, desmin and N-WASP-depletion and increased YAP in nuclei at the basal state along with an absence of any further response to cyclic stretching (Fig. 4C and Supplementary Fig. 3I). In order to compare the effect of a long-term siRNA-mediated depletion to an acute endocytosis inhibition on YAP distribution, were treated myotubes with a dynamin GTPase activity inhibitors which drastically block endocytosis (Dynole 34-2). This inhibition significantly decreased basal nuclear YAP levels and abolished responsiveness to stretch (Fig. 4D). The involvement of DNM2 on YAP translocation prompted us to analyze YAP nucleocytoplasmic distribution in primary myotubes from a CNM mouse model, HTZ KI-*Dnm2*^R465W^ mice. These myotubes displayed the same YAP shuttling defects as DNM2-depleted mouse myotubes, with abnormally increased YAP nuclear levels at the basal state (Fig. 4E) and no response to cyclic stretching. We next analyzed immortalized human myotubes and compared YAP distribution with human myotubes from a patient expressing the *DNM2* p.R369Q CNM mutation. These myotubes displayed the exact same YAP shuttling defects as DNM2-depleted mouse myotubes, with abnormally increased YAP nuclear levels at the basal state and no response to cyclic stretching (Supplementary Fig. 4 D-E), stressing the importance of DNM2 function in controlling YAP signaling.

**Fig. 4.**
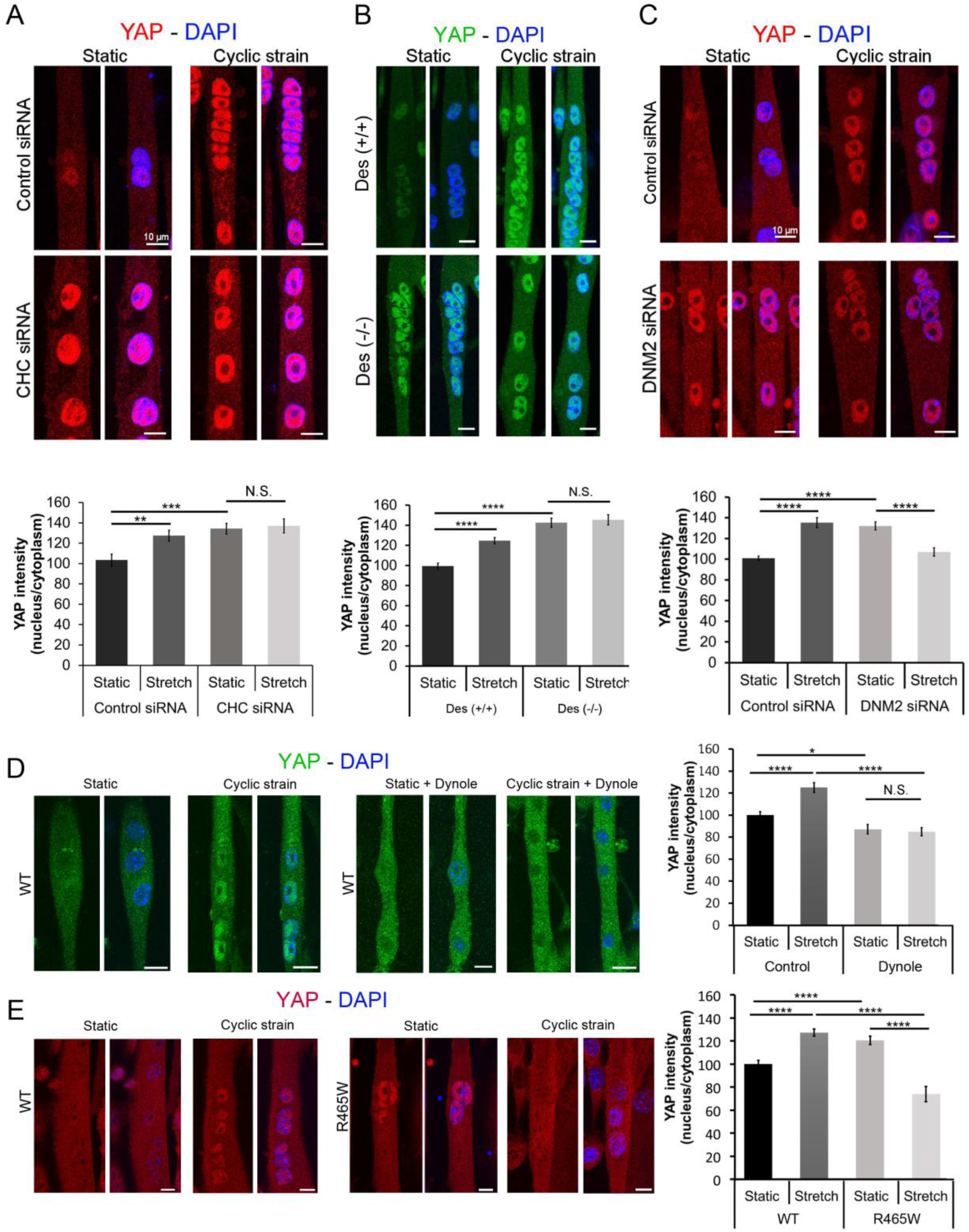
CHC, desmin and DNM2 depletion deregulate YAP signaling in differentiated myotubes. **(A)** Immunofluorescent staining of YAP (red) and nuclei (blue) in mouse primary control or CHC-depleted myotubes either stretched or left static. Bottom panel shows YAP fluorescence in control or CHC-depleted myotubes (*n* = 30–45 myotubes). **(B)** Immunofluorescent staining of YAP (red) and nuclei (blue) in mouse primary control or desmin knock-out myotubes (Des-/-) either stretched or left static. Bottom panel shows YAP fluorescence in control or Des-/-myotubes (*n* = 30–45 myotubes). **(C)** Immunofluorescent staining of YAP (red) and nuclei (blue) in mouse primary control or DNM2-depleted myotubes either stretched or left static. Bottom panel shows YAP fluorescence in control or DNM2-depleted myotubes (*n* = 30–45 myotubes). **(D)** YAP immunofluorescent staining (green) and nuclei (blue) in mouse primary myotubes treated with inactive negative control (31-2 compound) or with dynamin inhibitor dynole (34-2) and either stretched or left static. Right panel shows YAP fluorescence in mouse primary myotubes treated with or without dynamin inhibitor dynole (*n* = 18–35 myotubes). **(E)** YAP immunofluorescent staining (green) and nuclei (blue) in mouse primary control or DNM2^R465W^ (^R465W^) myotubes either stretched or left static. (*n* = 19–51 myotubes). Data are presented as mean ± SEM; *P < 0.05, **P < 0.01, ***P < 0.001, ****P < 0.0001.

### DNM2-linked CNM mutations disorganize clathrin plaques and desmin in vivo and delay plaque dynamics

The involvement of DNM2 mutants on YAP translocation prompted us to analyze clathrin plaques and the desmin network in *DNM2*-related CNM. We first analyzed clathrin plaques in a knock-in mouse model for the most frequent human mutation, i.e. heterozygous (HTZ) KI-*Dnm2*^R465W^ mice by intravital microscopy (Fig. 5A). Given that the AP2 adaptor is specifically recruited at the PM and is a *bona-fide* component of clathrin plaques in muscle (Vassilopoulos et al., 2014), it was used as marker to study the plaques *in vivo*. We transduced the tibialis anterior (TA) muscle with an adeno-associated virus (AAV9) expressing the µ2-subunit of the AP2 clathrin adaptor tagged with mCherry (AP2-mCherry). In WT mice, AP2-mCherry was expressed at the surface of muscle fibers, aligned with the Z-bands and colocalized with endogenous AP2, CHC, DNM2 and α-actinin 2 (Fig. 5B left and Supplementary Fig. 5A-D left). In HTZ KI-*Dnm2*^R465W^ mice, AP2 distribution was discontinuous and AP2 patches were completely disorganized in the most severely affected regions (Fig. 5C right and Supplementary Fig. 5A-D right).

**Fig. 5.**
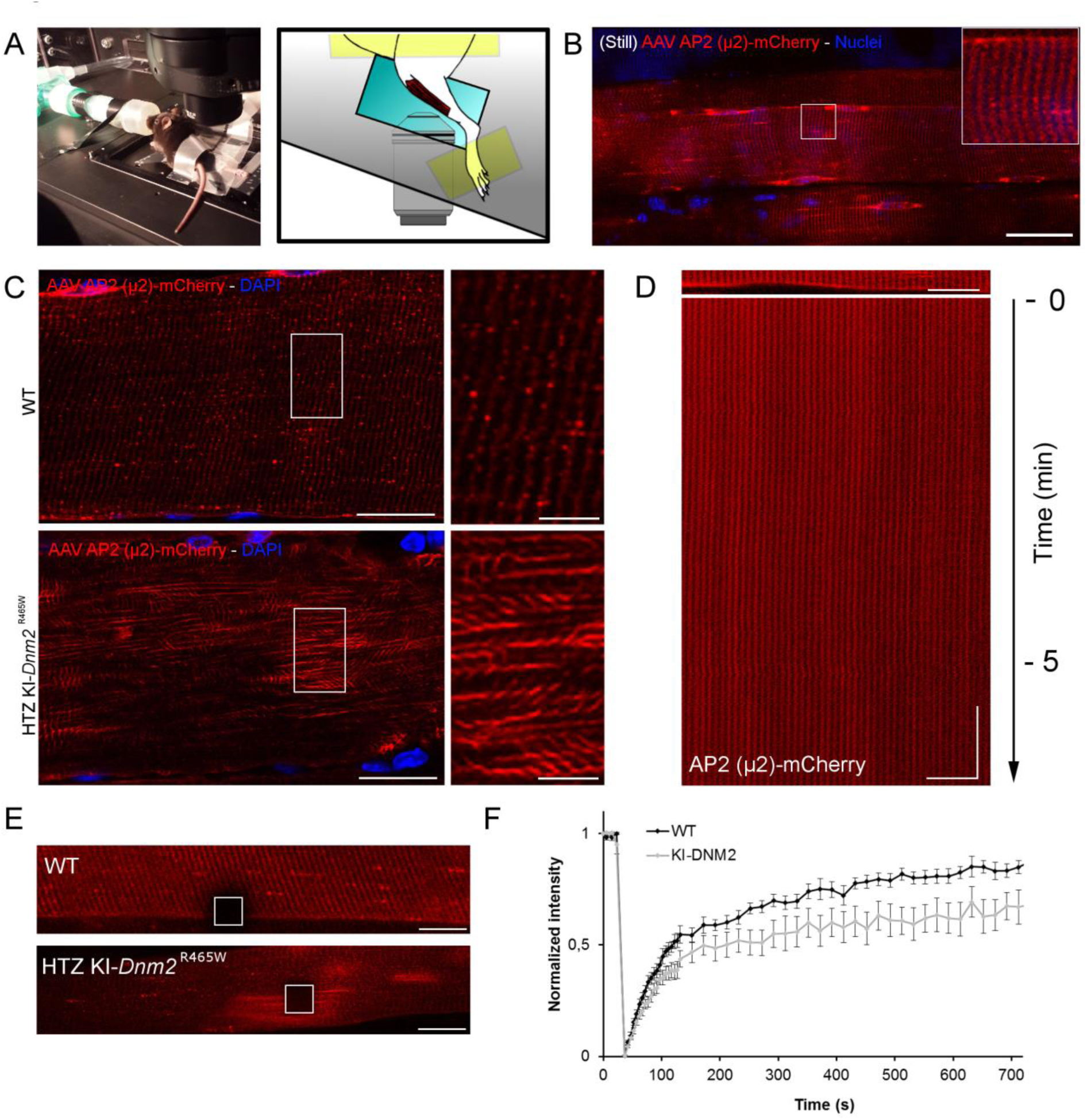
Intravital clathrin plaque imaging in WT and HTZ KI-*Dnm2* ^R465W^ mice. **(A)** Intravital set-up with live anesthetized mouse (left panel). Cartoon of the set-up used to image superficial TA fibers in live mice (right panel). **(B)** WT mouse muscle imaged *in vivo* using confocal microscopy. Note AP2 striation in inset. Bar is 50 µm. **(C)** AAV9-AP2 mcherry distribution between WT and HTZ KI-*Dnm2* ^R465W^ mice at the surface of fixed longitudinal muscle sections observed using confocal microscopy. Bars are 20 µm and 5 µm for insets. **(D)** Kymograph of mouse muscle injected with AAV9-AP2 mcherry imaged over 6 minute time-lapse Note that the signal is stable throughout the whole experiment. Horizontal bar is 20 µm and vertical bar is 1 min. **(E)** Representative FRAP regions (box) from mouse muscle injected with AAV9-AP2 mcherry. Bars are 20 µm. **(F)** Quantification of AP2-mcherry fluorescence recovery between WT and HTZ KI-*Dnm2* ^R465W^ mice. Two-WAY ANOVA was performed, ****P < 0.0001 (*n*=17 to 25 fibers from at least 3 mice for each genotype).

Our results on the abnormal distribution of AP2 in muscle from HTZ KI-*Dnm2*^R465W^ mice suggesting disorganized clathrin plaques prompted us to analyze their dynamics *in vivo*. In WT mice, the distribution of AP2 monitored on a kymograph was stable through time, even when imaged at fast acquisition frequencies for periods which exceed the average time that it takes to form an endocytic vesicle (Fig. 5D and Supplemental Movie 2). In addition, *in vivo* AP2 fluorescence recovery after photobleaching (FRAP) was dynamic (Supplemental Movie 3), reaching a 85% plateau in 10 minutes (Fig. 5E, F) in agreement with *in vitro* experiments (Wu et al., 2003). In HTZ KI-*Dnm2*^R465W^ mice, the speed of fluorescence recovery of AP2 was not significantly delayed, although the plateau only reached 60% (Fig. 5E, F and Supplemental Movie 4). This decrease in the mobile fraction of AP2 observed in HTZ KI-*Dnm2*^R465W^ mice suggests defective exchange of plaque components. Since DNM2 participates in cytoskeletal organization and flat clathrin plaque homeostasis *in vitro*, we reasoned that CNM-causing mutations could induce defects in plasma membrane IF organization and lead to malfunction of muscle costameres *in vivo*. We investigated the organization of the desmin network in dissociated fibers from HTZ KI-*Dnm2* ^R465W^ mice. Confocal sections from the top to the bottom of dissociated fibers revealed disorganization of desmin distribution at the surface of isolated fibers from HTZ KI-*Dnm2* ^R465W^ mice (Fig. 6A). Analysis of muscle from HTZ KI-*Dnm2* ^R465W^ mice using thin section EM revealed presence of large regions of the surface containing IF tangles (Fig. 6B-E), resembling those observed in CHC- and AP2-depleted cells and which were very pronounced in DNM2-depleted myotubes. These characteristic aggregates were also present around central nuclei in HTZ KI-*Dnm2* ^R465W^ mice, but not around the rare central nuclei of WT mice (Fig. 6F-H). The results from *DNM2* knock-in mice prompted us to investigate the organization of desmin IFs in CNM patients with *DNM2* mutations. Muscle biopsies from *DNM2*-related CNM patients present a characteristic radial arrangement of sarcoplasmic strands particularly visible with oxidative enzyme reactions. Transverse muscle sections from patients harboring either the p.^R465W^ or the p.R369Q DNM2 mutation presented a strong desmin accumulation in the radiating sarcoplasmic strands, typical of dilated Z-band material, which stretched from central nuclei towards the sarcolemma (Fig. 6I-M and Supplementary Fig. 6A). We also analyzed the distribution of YAP on muscle sections from CNM patients harboring either the p.R369Q, p.^R465W^ or an additional p.R369W DNM2 mutation. In control skeletal muscle YAP was localized in the cytoplasm, and no myonuclei were labelled (Supplementary Fig. 6B). Interestingly, CNM patients presented regions with increased nuclear YAP staining in both centralized and peripheral myonuclei.

**Fig. 6.**
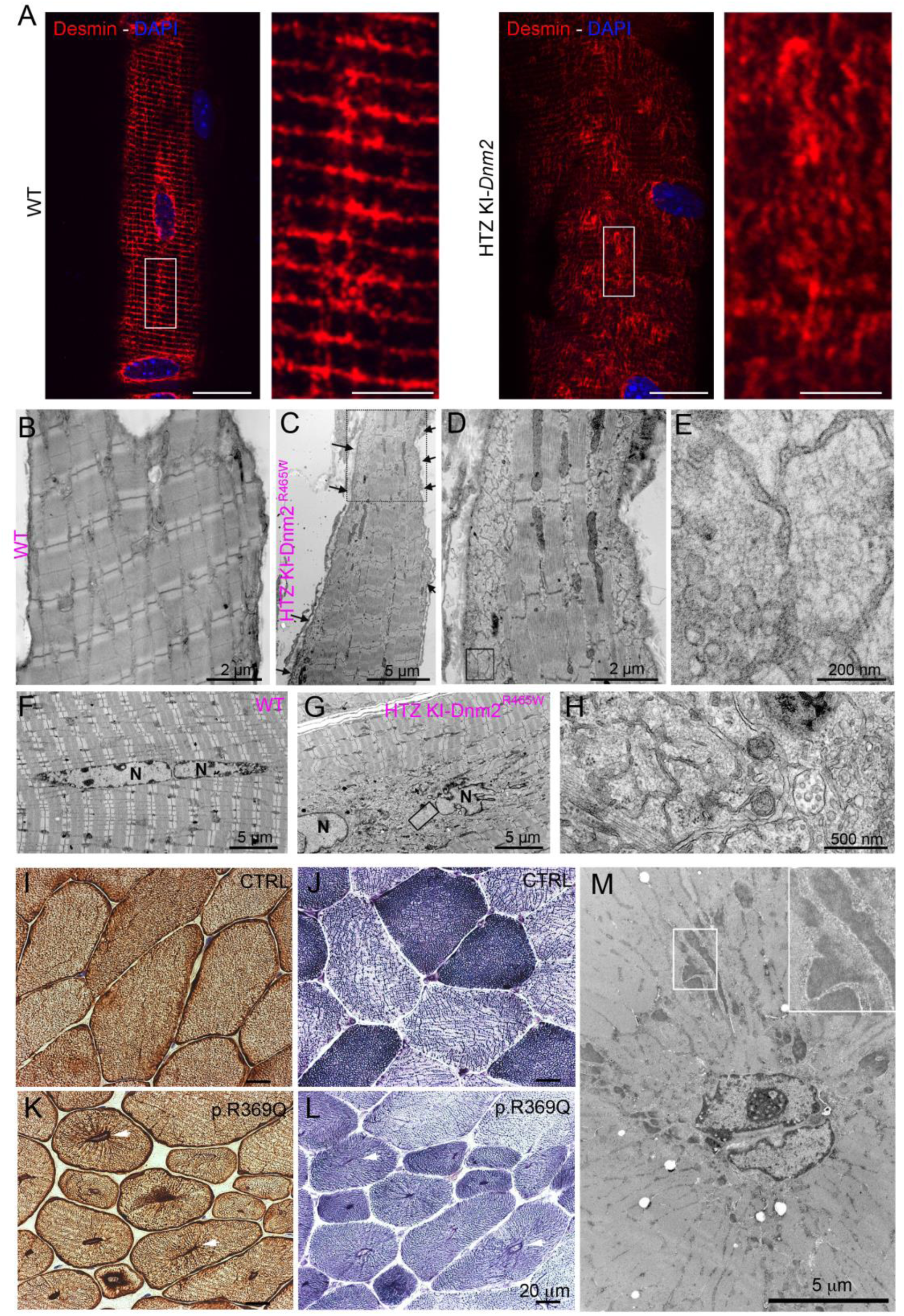
Desmin IF defects in HTZ KI-*Dnm2* ^R465W^ mice and human CNM patients. **(A)** Confocal sections from the surface of WT (left panel) or HTZ KI-*Dnm2* ^R465W^ (right panel) mouse dissociated skeletal muscle fibers immunolabeled with desmin antibody. Bars are 20 µm and 5 µm for insets. **(B-E)** Thin-section EM of surface longitudinal skeletal muscle sections from either WT (B) or HTZ KI-*Dnm2* ^R465W^ (C-E) mice. The presence of desmin filament aggregates containing ER membranes at the periphery of the fiber are clearly visible on the insets. **(F-H)** Longitudinal skeletal muscle sections from the core of either WT (F) or HTZ KI-*Dnm2* ^R465W^ mice (G-H) displaying central nuclei. The presence of desmin filament aggregates containing ER membranes around myonuclei is clearly visible in the inset (H). **(I-K)** Muscle biopsy sections of a control patient (I) and a patient with p.R369Q *DNM2 mutation* immunohistochemically labelled against desmin. Muscle sections from the patient reveal a strong desmin labelling of radial sarcoplasmic strands. **(J)** NADH-TR reaction C on healthy muscle section. **(L)** NADH-TR reaction C on muscle section from a CNM patient harboring a p.R369Q mutation display a high percentage of small rounded fibers with centralized nuclei and few fibers with typical aspect of radiating sarcoplasmic strands. **(M)** Thin-section EM of muscle from CNM patient with the p.^R465W^ mutation presenting characteristic radial strands. Note the characteristic dilated Z-band material in the inset.

## Discussion

Collectively, our experiments show the existence of a novel compartment centered on clathrin plaques and branched actin which lies at the crossroads of mechanotransduction and cytoskeleton-mediated mechanosensing. We show that actin filaments surrounding mechanically sensitive clathrin plaques form a scaffold for YAP at the membrane and are connected to other parts of the cell by a three-dimensional intermediate filament web. By virtue of shaping both clathrin lattices and branched actin filaments, DNM2 takes center stage as a regulator of YAP signaling and intermediate filaments anchoring.

Comparable to focal adhesions, clathrin plaques are macromolecular functional units that connect extracellular matrix, plasma membrane and intracellular cytoskeleton. Our work shows that branched actin filaments which form around adhesive clathrin plaques are the central element for both IF anchoring and YAP recruitment at the PM and we identified DNM2 and N-WASP as central actors of actin remodeling at these sites. We have previously shown that the actin network surrounding clathrin lattices contains α-actinin 2, an actin crosslinking protein (Vassilopoulos et al., 2014). The same actin structures require Arp2/3 activity including involvement of the actin nucleation-promoting factor N-WASP in Hela cells (Leyton-Puig et al., 2017). It is of interest that hybrid Arp2/3 and α-actinin containing complexes have already been reported (Chorev et al., 2014; Pizarro-Cerdá et al., 2016) and association of clathrin plaques with these tissue-specific hybrid complexes could be the signature of clathrin’s role in adhesion and mechanotransduction. DNM2, previously shown to associate with flat clathrin structures by EM (Damke et al., 1994; Sochacki et al., 2017; Warnock et al., 1997) interacts with the cellular machinery, including N-WASP, that induces branched actin polymerization around clathrin plaques. We also provide the first evidence that intermediate filaments of desmin, prominent in cells subject to mechanical stress and capable of connecting the cell surface with peripheral nuclei, are systematically associated with actin structures surrounding clathrin plaques. Depleting either CHC, AP2, N-WASP or DNM2 induced formation of IF aggregates such as those found in patients with desmin mutations (Clemen et al., 2015) demonstrating the strong dependence of the intermediate cytoskeleton organization to the clathrin plaques and associated machinery.

It has been established that in proliferating myoblasts, YAP/TAZ are localized in the nucleus where they act as coactivators for several transcription factors and that during differentiation, both translocate from the nucleus to the cytoplasm (Watt et al., 2010). In addition, mechanical cues such as cell stretching can cause YAP/TAZ translocation back into the nucleus (Cui et al., 2015; Fischer et al., 2016). We show that clathrin plaques and associated cytoskeletons might serve as signaling platforms that respond to mechanical cues triggering YAP signaling. Metal-replica EM analysis allowed us to map YAP molecules on actin filaments surrounding clathrin-coated structures suggesting that the same branched actin structures which anchor desmin IFs are also involved in YAP scaffolding. Upon cyclic stretching mimicking muscle contraction clathrin plaque size was decreased along with an increased clathrin-mediated endocytosis. This increased endocytosis must occur during the relaxation phases as it has been shown that endocytosis efficiency is inversely correlated to higher membrane tension (Boulant et al., 2011). Our results demonstrate that upon stretching, the plaque size is drastically reduced along a concomitant YAP increase in the nuclei of differentiated myotubes. Taken together, these data open the possibility that during stretching, clathrin plaque remodeling by endocytosis releases a pool of YAP necessary for its nuclear translocation. In agreement, depletion of clathrin, DNM2 and N-WASP over a relatively long period of time resulted in increased nuclear YAP at the basal level of differentiated myotubes while acute dynamin GTPase inhibition blunted YAP nuclear translocation upon stretching suggesting a contribution of clathrin-mediated endocytosis in this process. This observation is in agreement with the presence of YAP directly on the actin filaments surrounding canonical clathrin-coated pits. Since branched actin filaments involved in clathrin-mediated endocytosis depolymerize once the fission of the budding vesicle is completed (Taylor et al., 2011), it is conceivable that YAP molecules could be released as actin filaments would disassemble. The same would apply to clathrin-coated pits budding from plaques. *DNM2* mutations, causing the autosomal dominant CNM, mostly segregate in the middle domain involved in DNM2 oligomerization and actin remodeling (Durieux et al., 2010; González-Jamett et al., 2017). This domain has been shown to directly interact with actin, and it is highly probable that the CNM phenotype observed in patients with DNM2 mutations directly stems at least in part from defective cortical actin turnover. These defects would produce dysfunctional force transmission at costameres explaining atrophy and reduced force generation in muscle from the KI-*Dnm2*^R465W^ mouse model (Durieux et al., 2010). Mechanistic understanding of CNM pathophysiology stemming directly from this work concerns the interplay between nuclear positioning and the IF tangles described here. The link between desmin IFs with both PM and peripheral nuclei in muscle is well established and it was recently shown that Arp2/3 and actin organize desmin for peripheral nuclear positioning (Roman et al., 2017). Our results suggest the peculiar organization termed "radial sarcoplasmic strands", found in CNM patient fibers, is a consequence of the desmin tangles systematically forming at the surface and around nuclei. The desmin tangles could adopt this radiating appearance characteristic of this disease during nuclear centralization. Using the *DNM2*-related CNM knock-in mouse model and patient myotubes we show increased basal nuclear YAP corroborating the effect produced by DNM2 depletion and strengthening the link between YAP signaling and DNM2 function. It is noteworthy that transgenic mice constitutively expressing nuclear YAP in skeletal muscle display a CNM-like phenotype (Judson et al., 2013). Given that desmin defects have been previously identified in the X-linked recessive form of CNM (Hnia et al., 2011), we may hypothesize that desmin IFs and YAP signaling defects could be a common component of CNM pathophysiology. Further investigation for desmin and YAP signaling defects in the autosomal recessive form of CNM due to amphiphysin 2 mutations (Nicot et al., 2007) will be warranted to confirm this common pathophysiological pathway. Importantly, the clathrin plaque-based structure may be the Achilles’ heel of several tissues and its dysfunction may lead to additional disorders including cancer where abnormal clathrin plaque assembly or misregulated DNM2 function could perturb the fine coupling between adhesion and force transduction.

## Materials and methods

### Antibodies

Primary antibodies are listed in Supplementary Table 2. Secondary antibodies for immunofluorescence were AlexaFluor-488, AlexaFluor-568 and AlexaFluor-647 conjugates (Life Technologies, France). Secondary antibodies were coupled to horseradish peroxidase (Jackson Laboratories, USA). Secondary antibodies used after immunoprecipitation were Trueblot IgG HRP from Rockland Inc., USA.

### Human myoblast cultures

Immortalized control and patient myoblast cells (DNM2 p.R369Q) were cultured in proliferation medium (1 volume of M199, #41150020, Invitrogen, France), 4 volumes of DMEM Glutamax, 20% fetal bovine serum (FBS), 50 U/mL penicillin, 50 mg/mL streptomycin, 25 μg/mL fetuin (#10344026; Life Technologies, France), 0.5 ng/mL fibroblast growth factor-basic (bFGF; #PHG0026; Life Technologies, France), 5 ng/mL human epidermal growth factor (hEGF; #PHG0311; Life Technologies, France), 0.2 μg/mL dexamethasone (#D4902-100mg; Sigma, France), 5 μg/mL insulin (#91077C-1g; Sigma, France). Differentiation was induced using DMEM Glutamax, 2% Horse Serum, 50 U/mL penicillin, 50 mg/mL streptomycin, supplemented with 5µg/mL insulin (#91077C-1g; Sigma, France).

### Mouse myoblast cultures and siRNA-mediated knock-down

Primary skeletal muscle cells were prepared from 3-4 day-old mouse pups. Cells were maintained in tissue culture dishes coated with Matrigel Matrix (Corning, France) in basal medium with 20% FBS, 50 U/mL penicillin, 50 mg/mL streptomycin (growth medium) and 1% Chicken Embryo Extract (Seralab, UK). Differentiation was induced when cells were ~80% confluent by switching to differentiation medium (basal medium with 2% horse serum). For experiments in Figure 1, in order to avoid detachment due to strong contractions, and to keep cells in culture for prolonged periods of differentiation, myotubes were covered with a layer of Matrigel Growth Factor Reduced (GFR) Basement Membrane Matrix, Phenol Red-Free (Corning, France) (Falcone et al., 2014). For siRNA treatment, cells (differentiated for either 4 or 6 days) were transfected twice for 48 hours using 200 nM siRNA and HiPerfect transfection reagent (Qiagen, Germany) according to manufacturer’s instructions. Targeting and control siRNAs were synthesized by Eurogentec, Belgium. The list of siRNAs used and sequences can be found in Supplementary Table 1. For CHC, AP2 and DNM2, sequences targeted were already published (Ezratty et al., 2009; Vassilopoulos et al., 2014).

### Electron microscopy of unroofed myotubes

Adherent PM from cultured cells grown on glass coverslips were obtained by sonication as described previously (Heuser, 2000). Sample processing for platinum-replica electron microscopy of unroofed cells was performed as follows: 2% glutaraldehyde/ 2% paraformaldehyde-fixed cells were further sequentially treated with 1% OsO4, 1% tannic acid and 1% uranyl acetate prior to graded ethanol dehydration and Hexamethyldisilazane (HMDS) (Sigma-Aldrich, France). For immunogold labeling, 4% paraformaldehyde fixed PMs were washed and quenched before incubation with primary and 15 nm gold-coupled secondary antibodies and further fixed with 2% glutaraldehyde. Dried samples were then rotary-shadowed with 2 nm of platinum and 5-8 nm of carbon using an ACE600 metal coater (Leica Microsystems, Germany). The resultant platinum replica was floated off the glass with hydrofluoric acid (5%), washed several times on distilled water, and picked up on 200 mesh formvar/carbon-coated EM grids. The grids were mounted in a eucentric side-entry goniometer stage of a transmission electron microscope operated at 80 kV (model CM120; Philips) and images were recorded with a Morada digital camera (Olympus, Tokyo). Images were processed in Adobe Photoshop to adjust brightness and contrast and presented in inverted contrast. Anaglyphs were made by converting the −10° tilt image to red and the +10° tilt image to cyan (blue/green), layering them on top of each other using the screen blending mode in Adobe Photoshop, and aligning them to each other. Tomograms were made by collecting images at the tilt angles up to ±25° relative to the plane of the sample with 5° increments. Images were aligned by layering them on top of each other in Adobe Photoshop.

### Dissociated fibers

Myofibers were isolated by mechanical dissociation from the dissected *tibialis anterior* (TA) muscle of 3-month old or 7-month old mice fixed 48 hours in 4% paraformaldehyde.

### Immunofluorescence microscopy

Adult mouse skeletal muscle was embedded in Tissue-Tek OCT compound (Miles Inc.), frozen, and stored at −80°C. Cryosections (10 µm thick) were fixed (15 min, 4% paraformaldehyde in PBS) at room temperature (RT), permeabilized (10 min, 0.5% Triton X-100 in PBS, RT) and blocked (30 min, PBS with 0.1% Triton X-100, 5% bovine serum albumin (BSA). Sections were incubated with primary antibodies (overnight, 4°C, in PBS with 0.1% Triton X-100, 5% BSA) and washed in PBS with 0.1% Triton X-100. Sections were then incubated with secondary antibodies (60 min, RT), washed in PBS with 0.1% Triton X-100, and mounted with Vectashield anti-fading solution containing DAPI (Vector Laboratories, USA). For double or triple labeling, the primary antibodies (from different species) were added simultaneously at the appropriate step.

For mouse and human cells grown on coverslips, cells were washed in warm PBS, fixed in paraformaldehyde (4% in PBS, 15 min), then washed in PBS, permeabilized (10 min, 0.5% Triton X-100 in PBS) and blocked (5% BSA in PBS with 0.1% Triton X-100, 30 min). Antibody labeling was performed by addition of 200 µL blocking solution with primary or secondary antibodies and washing with PBS with 0.1% Triton X-100. F-Actin was stained using Alexa Fluor 555 Phalloidin and G-actin was stained using Alexa 488 DNase I (Thermo Fisher Scientific, France) for 1h at RT. Samples were mounted in Vectashield containing DAPI (Vector Laboratories, USA).

Samples were analyzed by confocal laser scanning microscopy using an upright FV-1000 confocal laser scanning microscope (Olympus, Tokyo) equipped with UPlanS-Apo 60x, 1.35 NA oil immersion objective lenses or a SP5 inversed microscope (Leica, Germany) equipped with a Leica HyD hybrid detector. Super resolution spinning-disk confocal images presented in Fig. 3A were taken with a Nikon Ti2 microscope equipped with a motorized stage and a Yokogawa CSU-W1 spinning disk head coupled with a Prime 95 sCMOS camera (Photometrics). To obtain super resolutive images, a Live SR module (Roper) was used. DAPI, Alexa-488 and Alexa-568 were sequentially excited using lasers with wavelengths of 405 for DAPI, 473 for Alexa-488 and 543 nm for Alexa-568. Z-series from the top to the bottom of fibers were sequentially collected for each channel with a step of 0.8-1µm between each frame. Imaging was performed at RT using Leica Type F immersion oil. Images (1024 × 1024 pixels) were saved as TIFF files in OLYMPUS FV-1000 software, and levels were adjusted in Adobe Photoshop or Gimp software. Image quantification was performed using National Institutes of Health’s FIJI (Schindelin et al., 2012).

### Cyclic stretch

Cells were plated onto flexible-bottom UniFlex or BioFlex plates (Flexcell International, USA) coated with Corning Matrigel GFR Basement Membrane Matrix and incubated at 37°C in a CO_2_ incubator. The cells were subjected to cyclic stretch at 0.5 Hz during 6 hours using a computer-controlled vacuum FX-4000T Tension Plus System stretch apparatus (FlexCell International, USA) with a vacuum pressure that is sufficient to generate 10% mechanical stretch. Replicate control samples were maintained under static conditions with no applied cyclic stretch. After stretching, cells were washed with PBS and fixed using 4% paraformaldehyde for immunofluorescence or NaCl-EDTA buffer for western blotting. For dynamin inhibition, myotubes were treated with or without 10 µM Dynole 34-2 or the inactive compound 31-2 as a negative control (Abcam, France).

### Transferrin uptake assay

Cells were subjected to cyclic stretching for 4 hours before incubation for 15 min with AlexaFluor-488 fluorescently labeled human transferrin (40 μg/mL) (Molecular Probes, Life Technologies, France) concomitant to stretching. Endocytosis of this compound was stopped with ice-cold PBS washing and fixed using 4% paraformaldehyde.

Entire myotubes were imaged using stacks of 500 nm step, with an FV-1200 confocal microscope (Olympus, Tokyo) and 40x oil objective.

### Immunoblot analysis

Cell samples were collected using Laemli blue 4X directly or a NaCl (150 mM)-EDTA (10 mM) buffer with added proteinase inhibitor cocktail (Sigma-Aldrich, France).

Protein samples were separated by electrophoresis (4-12% bis-acrylamide gel, Life Technologies, France), then electrophoretically transferred to 0.45 µm nitrocellulose membranes (Life Technologies, France) and labelled with primary antibodies and secondary antibodies coupled to horseradish peroxidase. The presence of proteins in samples was detected using Immobilon Western Chemiluminescent HRP Substrate (Sigma-Alrich, France). Acquisition was performed on a G-Box (Ozyme, France).

### Immunoprecipitation

IPs were performed on primary cell cultures. Myotube pellets were resuspended in 500 µL of lysis buffer (50 mM Tris-HCl, pH 7.5, 0.15 M NaCl, 1 mM EDTA, 1% NP-40) and protein inhibitor cocktail 1:100 (Sigma Aldrich, France). Each sample (200-500 µg) was precleared twice with 30 µL Protein-G-Sepharose (PGS 4 fast flow, Thermo Fisher, France) and incubated with 20 µg of specific antibody overnight (4°C). Washed PGS (40 µL) was first incubated with BSA (2 g/L) and further incubated with samples for 2 hours at 4°C. Pelleted PGS was taken up in sample buffer and subjected to electrophoresis and immunoblotting. For all immunoprecipitation experiments, HRP-conjugated rabbit and mouse IgG TrueBlot secondary antibodies (Rockland Inc., USA) were used.

### Intravital-imaging

WT and HTZ KI-*Dnm2*^R465W^ mice (Durieux et al., 2010) injected a month prior with AAV-µ2-mCherry were anesthetized using isofluorane. Skin was removed from the TA before applying muscle directly on coverslip and immobilizing with tape (Figure 5A). Imaging and FRAP was performed on an inverted Leica SP8 multiphoton microscope optimized for intravital microscopy of small animals, equipped with a complete isofluorane anesthesia unit. The mouse temperature was maintained at 37°C with a heating stage and a heating lamp, and breathing of the animal was monitored visually throughout the experiment.

### AAV production, titration and *in vivo* gene transfer

An adeno-associated virus serotype 9 was produced for expression of the µ2-subunit of the AP2 clathrin adaptor tagged with mCherry (AP2-mCherry) (Taylor et al., 2011). AAV2/9 pseudotyped vectors were prepared by tri-transfection in 293 cells as described previously (Rivière et al., 2006) using the pSMD2-AP2-mCherry plasmid, pXX6 plasmid coding for the viral sequences essential for AAV production and the pRepCap plasmid coding for serotype 9 capsid. Vector particles were purified on iodixanol gradient and concentrated on Amicon Ultra-15 100K columns (Merck-Millipore, USA). The viral genomes titer (vg/mL) was determined by quantitative real-time PCR. WT and HTZ KI-Dnm2^R465W^ mice were injected at 6 months of age. One intramuscular injection (40 µL/TA) of AAV9-AP2-mCherry was performed in TA muscles using 29G needle.

### Histomorphological and ultrastructural analyses

Human open muscle biopsies from two patients carrying the CNM-dynamin-2 mutation p.^R465W^, one patient carrying the CNM-dynamin-2 mutation p.R369Q, one patient carrying the CNM-dynamin-2 mutation p.R369W and two healthy control muscle were performed at the Centre de Référence de Pathologie Neuromusculaire Paris-Est, Institut de Myologie, GHU Pitié-Salpêtrière, Assistance Publique-Hôpitaux de Paris, GH Pitié-Salpêtrière, Paris, France, following written informed consent specially dedicated for diagnosis and research. Muscle was frozen in liquid nitrogen-cooled isopentane. For conventional histochemical techniques on human biopsies, 10 μm thick cryostat sections were stained with antibodies against desmin, YAP or with reduced nicotinamide adenine dinucleotide dehydrogenase-tetrazolium reductase (NADH-TR) by standard methods. Pictures of each section were obtained with a Zeiss AxioCam HRc linked to a Zeiss Axioplan Bright Field Microscope and processed with the Axio Vision 4.4 software (Zeiss, Germany).

For thin-section EM, mouse muscles were fixed by intra-aortic perfusion with 2% paraformaldehyde, 2% glutaraldehyde in 0.1M phosphate buffer (pH 7.4). Tibialis anterior samples were postfixed with 2% OsO4, in 0.1 M phosphate buffer (pH 7.4) for 1 h, then dehydrated in a graded series of acetone including a 1% uranyl acetate staining step in 70% acetone, and finally embedded in epoxy resin (EMBed-812, Electron Microscopy Sciences, USA). Myotubes grown on Thermanox coverslips (Nunc, Rochester, USA) were directly fixed for 30 min in the same fixation solution and processed as previously indicated for Tibialis anterior muscles. Ultra-thin (70 nm) sections were stained with uranyl acetate and lead citrate. For patient biopsies, frozen muscle sections (40 μm) were fixed in osmium tetroxide (1%), dehydrated and embedded in epoxy resin (as above). Ultra-thin (80 nm) sections were stained with uranyl acetate and lead citrate. Observations were made on a Philips CM120 electron microscope operated at 80kV (Philips, Eindhoven, The Netherlands) and images were recorded with a Morada digital camera (Olympus Soft Imaging Solutions GmbH, Münster, Germany).

### Image analysis

Desmin aggregate size analysis: the “Analyze particles” FIJI plugin (version 1.46) was used to count intracellular particles on binary confocal images of primary mouse myotubes in a single image from the middle of the cell. Same treatment was performed to measure the size of clathrin plaques on confocal images. Transferrin-AlexaFluor488 fluorescence was measured on confocal images by selecting 5 ROIs in myotubes, background noise, and using this formula: CTCF = Integrated density – (Area of selected cell × Mean fluorescence of background reading). FRAP analysis: Drifting was corrected using FIJI plugin JavaSIFT. Intensity of bleached area was compared to overall intensity of the muscle fiber, frame by frame, using FRAP Profiler plugin for FIJI.

### Statistics

Statistical analysis was performed using Student’s t-test except as otherwise stated.

### Study approval

Animal studies conform to the French laws and regulations concerning the use of animals for research and were approved by an external Ethical committee (approval n°00351.02 delivered by the French Ministry of Higher Education and Scientific Research). For human studies, all individuals provided informed consent for muscle biopsies according to a protocol approved by the ethics committee of the Centre de Référence de Pathologie Neuromusculaire Paris-Est, Institut de Myologie, GHU Pitié-Salpêtrière, Assistance Publique-Hôpitaux de Paris, GH Pitié-Salpêtrière, Paris, France.

### Data availability

All data supporting the findings of this study are available from the corresponding author on request.

### Author contributions

A.Fr., J.L. and M.B., designed and performed experiments, analyzed results and wrote the manuscript. G.M., M.T., C.G., A.Fo., S.B-Z., A.B., E.L., M.T.B., G.B. S.C. performed experiments. G.M., V.M., P.G., N.R., and C.C. analyzed the data. S.V supervised the study, designed and performed experiments and wrote the manuscript. All authors read and approved the final version of the manuscript.

## Acknowledgements

We are grateful to John Heuser and Azumi Yoshimura for precious help throughout this work. We dedicate this work to the memory of our friend Christien Merrifield. We also thank Onnik Agbulut and Athanassia Sotiropoulos for reagents, advices and comments, the Penn Vector Core, Gene Therapy Program (University of Pennsylvania, Philadelphia, US) for providing the plasmids for AAV construction, Laura Julien for AAV production, the Pitié-Salpêtrière (PICPS) and the Gustave Roussy institute imaging platforms for confocal and two-photon imaging facilities respectively and the IBPS electron microscopy platform. This work was supported by the Institut National de la Santé et de la Recherche Médicale (INSERM), Association Institut de Myologie (AIM), Sorbonne Université (SU), an Agence Nationale de la Recherche grant (ANR-14-CE12-0009 Dynamuscle) to M.B. and a young researcher grant (ANR-14-CE12-0001-01 Endomechano) to S.V.

---

**Supplementary Table 1.**
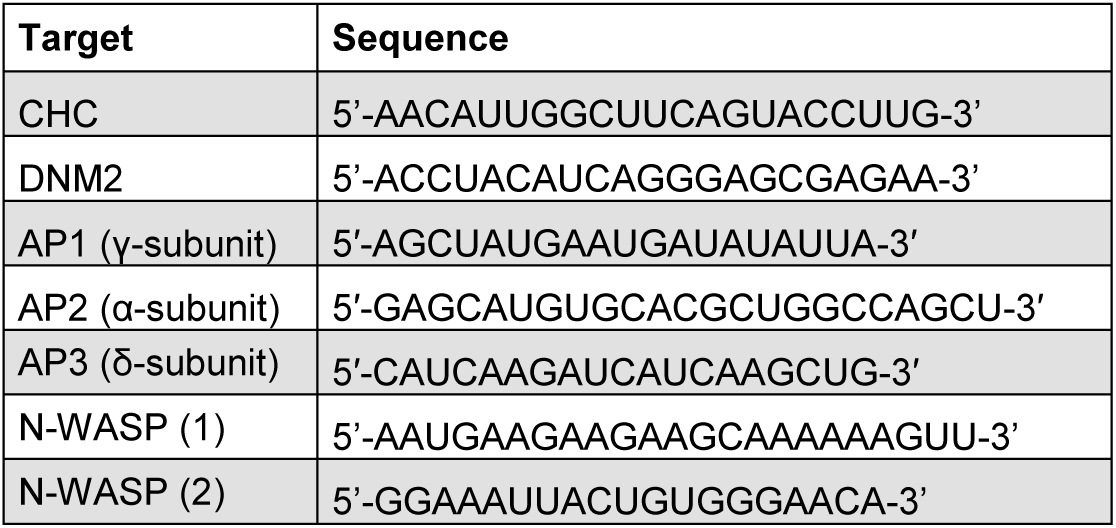
List of siRNA sequences

**Supplementary Table 2.**
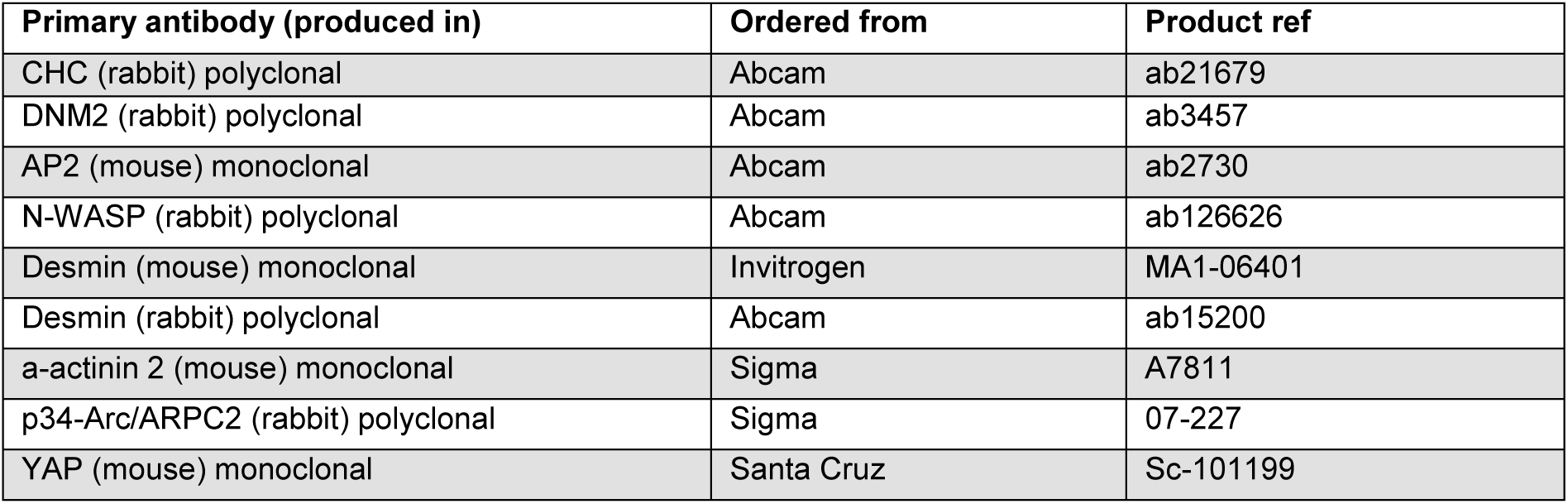
List of antibodies

| Primary antibody (produced in) | Ordered from | Product ref |
| --- | --- | --- |
| CHC (rabbit) polyclonal | Abcam | ab21679 |
| DNM2 (rabbit) polyclonal | Abcam | ab3457 |
| AP2 (mouse) monoclonal | Abcam | ab2730 |
| N-WASP (rabbit) polyclonal | Abcam | ab126626 |
| Desmin (mouse) monoclonal | Invitrogen | MA1-06401 |
| Desmin (rabbit) polyclonal | Abcam | ab15200 |
| α-actinin 2 (mouse) monoclonal | Sigma | A7811 |
| p34-Arc/ARPC2 (rabbit) polyclonal | Sigma | 07-227 |
| YAP (mouse) monoclonal | Santa Cruz | Sc-101199 |

**Supplementary Movie 1. Clathrin structures on the surface of muscle fibers anchor intermediate filaments.** Platinum replica EM imaging with electron tomography was achieved by collecting images at different tilt angles up to ±25° relative to the plane of the sample (Supplemental Movies S1). Clathrin lattices are pseudocolored in purple, branched actin in orange and intermediate filaments in green. Image size is 2198 × 885 nm.

**Supplementary Movie 2. Persistent clathrin structures on the surface of muscle fibers** Intravital imaging of TA muscle from WT mouse injected with AAV9-AP2 mCherry. Images were captured on a Leica SP8 multiphoton microscope at a frame rate of an image every 690 milliseconds over a 5 min time course. Bar 20 µm.

**Supplementary Movie 3. Recovery of AP2 fluorescence in WT mouse.** Intravital imaging of TA muscle from WT mouse injected with AAV9-AP2 mCherry. FRAP was performed after two frames and recovery of fluorescence was recorded several minutes after photobleaching. Time is shown in seconds.

**Supplementary Movie 4. Recovery of AP2 fluorescence in HTZ KI-Dnm2^R465W^ mouse.** Intravital imaging of TA muscle from HTZ KI-Dnm2^R465W^ mouse injected with AAV9-AP2 mCherry. FRAP was performed after two frames and recovery of fluorescence was recorded several minutes after photobleaching. Note the abnormal AP2 striations in knock-in mice. Time is shown in seconds.

## Supplementary material

**Supplementary Fig. 1.**
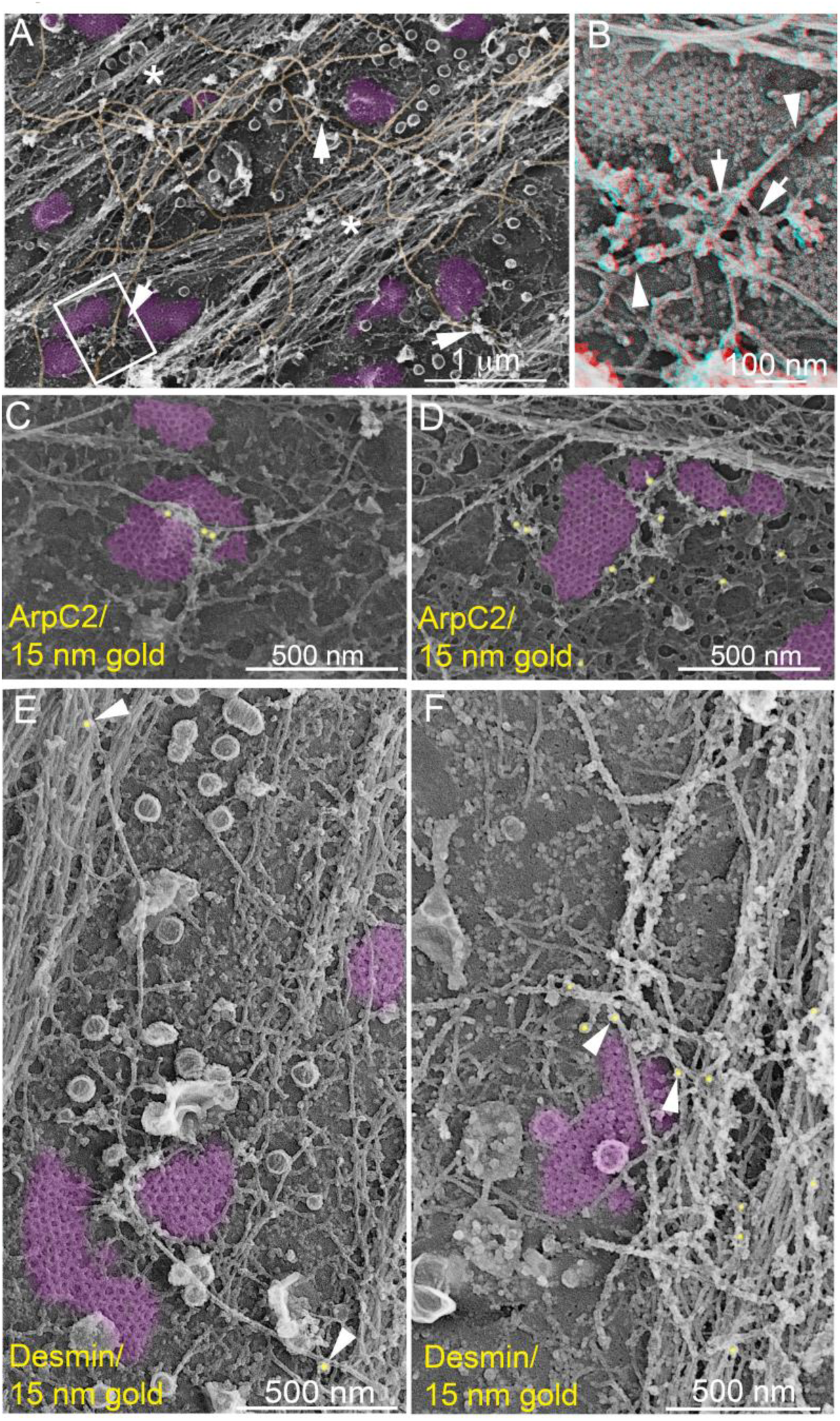
Clathrin lattices anchor desmin intermediate filaments and are surrounded by Arp2/3 branched actin filaments. **(A)** Survey view of unroofed C2C12 myotubes. Clathrin lattices are highlighted in purple and intermediate filaments in orange. Actin structures around clathrin plaques are indicated with arrows and asterisks denote typical stress fibers. **(B)** 3D anaglyph of the boxed region in (A). Use view glasses for 3D viewing of anaglyphs (left eye = red). Actin network is shown with arrows, while intermediate filaments are indicated with arrowheads. **(C) (D)** Branched actin surrounding clathrin plaques is immunolabeled using a primary antibody against ArpC2 and secondary antibodies coupled to 15 nm gold beads. Gold beads are pseudocolored yellow, clathrin lattices are highlighted in purple. **(E) (F)** Intermediate filaments are immunolabeled using a primary antibody against desmin and secondary antibodies coupled to 15 nm gold beads. Gold beads are pseudocolored yellow and indicated with arrowheads. Clathrin-coated structures are highlighted in purple.

**Supplementary Fig. 2.**
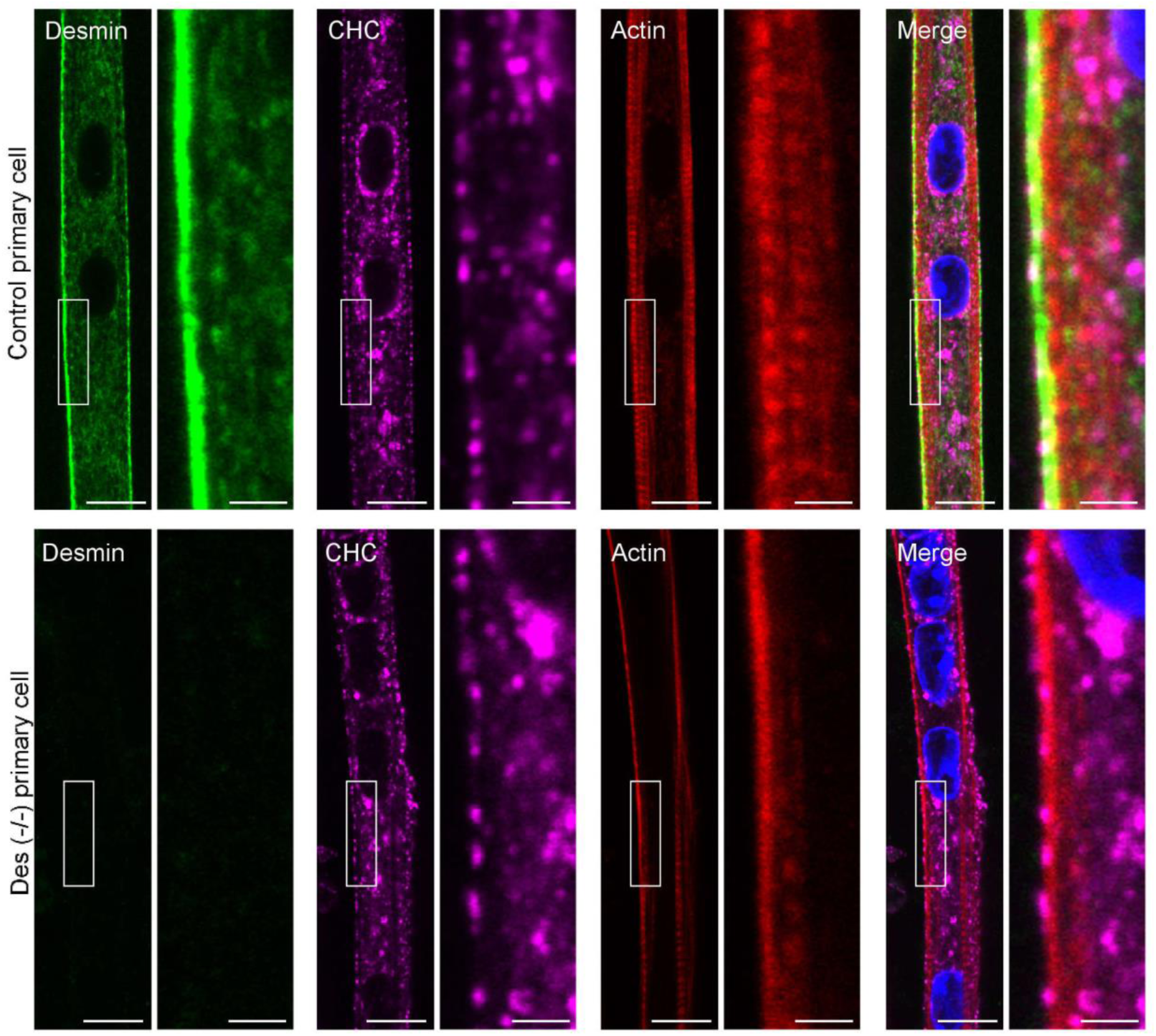
Clathrin plaque presence and increased basal nuclear YAP levels in desmin knock-out mice. Immunofluorescent detection ofdesmin (green), CHC (magenta) and actin staining (Phalloïdin, red) in differentiated mouse primary myotubes from WT mice (top panel) or myotubes from desmin knock-out mice (Des-/-, bottom panel). Bars 10 µm and 2 µm for insets.

**Supplementary Fig. 3.**
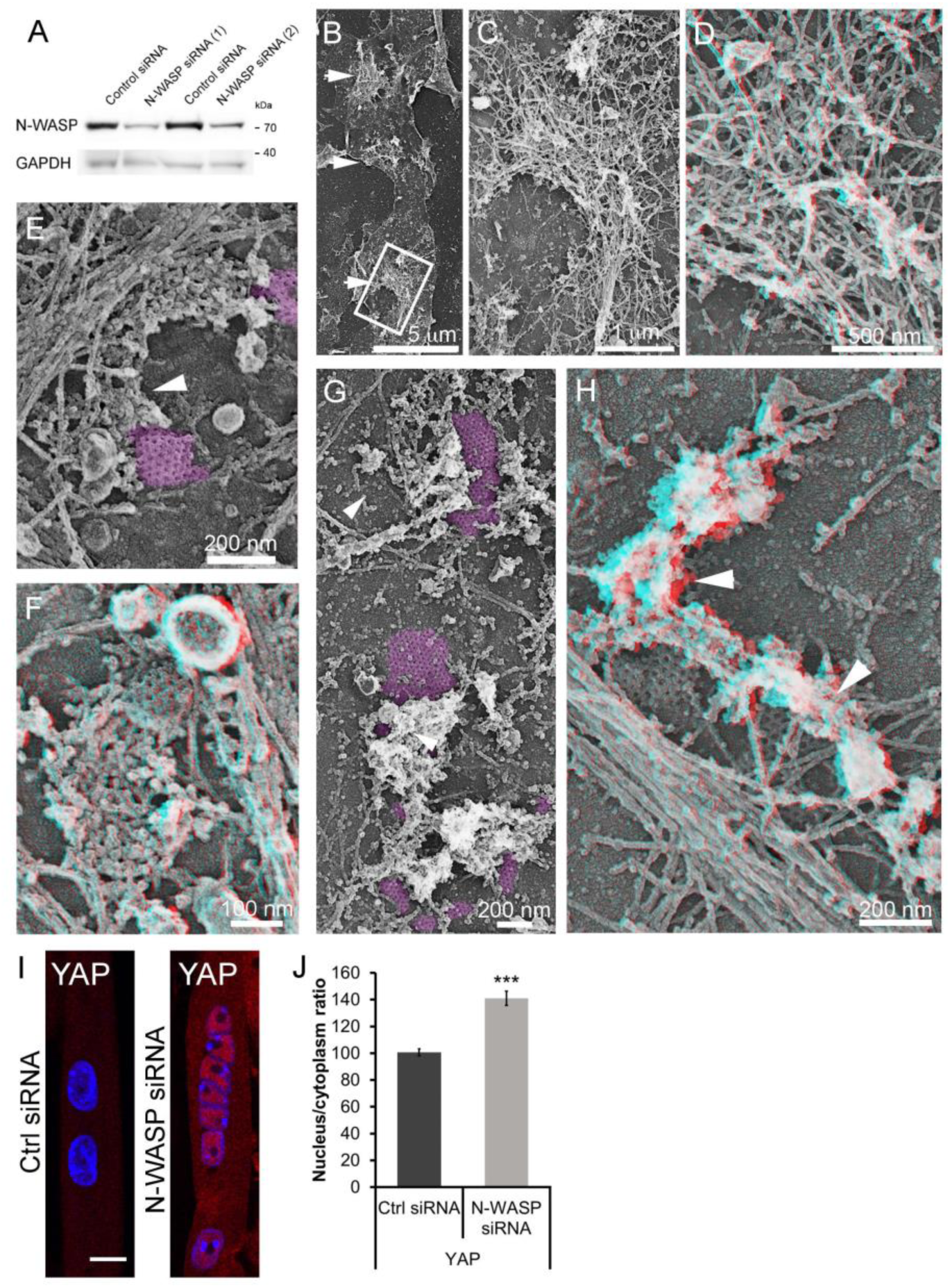
Desmin aggregates, abnormal actin around clathrin plaques and increased nuclear YAP basal levels in N-WASP depleted myotubes. **(A)** Myotubes were treated with control siRNA or two different siRNA targeting N-WASP and cell lysates immunoblotted for proteins (indicated left). **(B)** Survey view of the cytoplasmic surface of the plasma membrane in unroofed primary mouse myotubes treated with siRNA against N-WASP. Arrows indicate the presence of intermediate filament tangles still associated with the PM. **(C) (D)** Higher magnification view of IF tangles associated with the plasma membrane in unroofed primary mouse myotubes treated with siRNA against N-WASP. **(E-H)** Representative branched actin (arrowheads) around clathrin-coated structures (purple) in unroofed primary mouse myotubes treated with siRNA against N-WASP. **(I)** Immunofluorescent staining of YAP (red) and nuclei (blue) in mouse primary control or N-WASP depleted myotubes. Nuclei are stained with DAPI. **(J)** YAP fluorescence in control or N-WASP depleted myotubes (*n* = 30 myotubes). For (D), (F) and (H) use view glasses for 3D viewing of anaglyphs (left eye = red).

**Supplementary Fig. 4.**
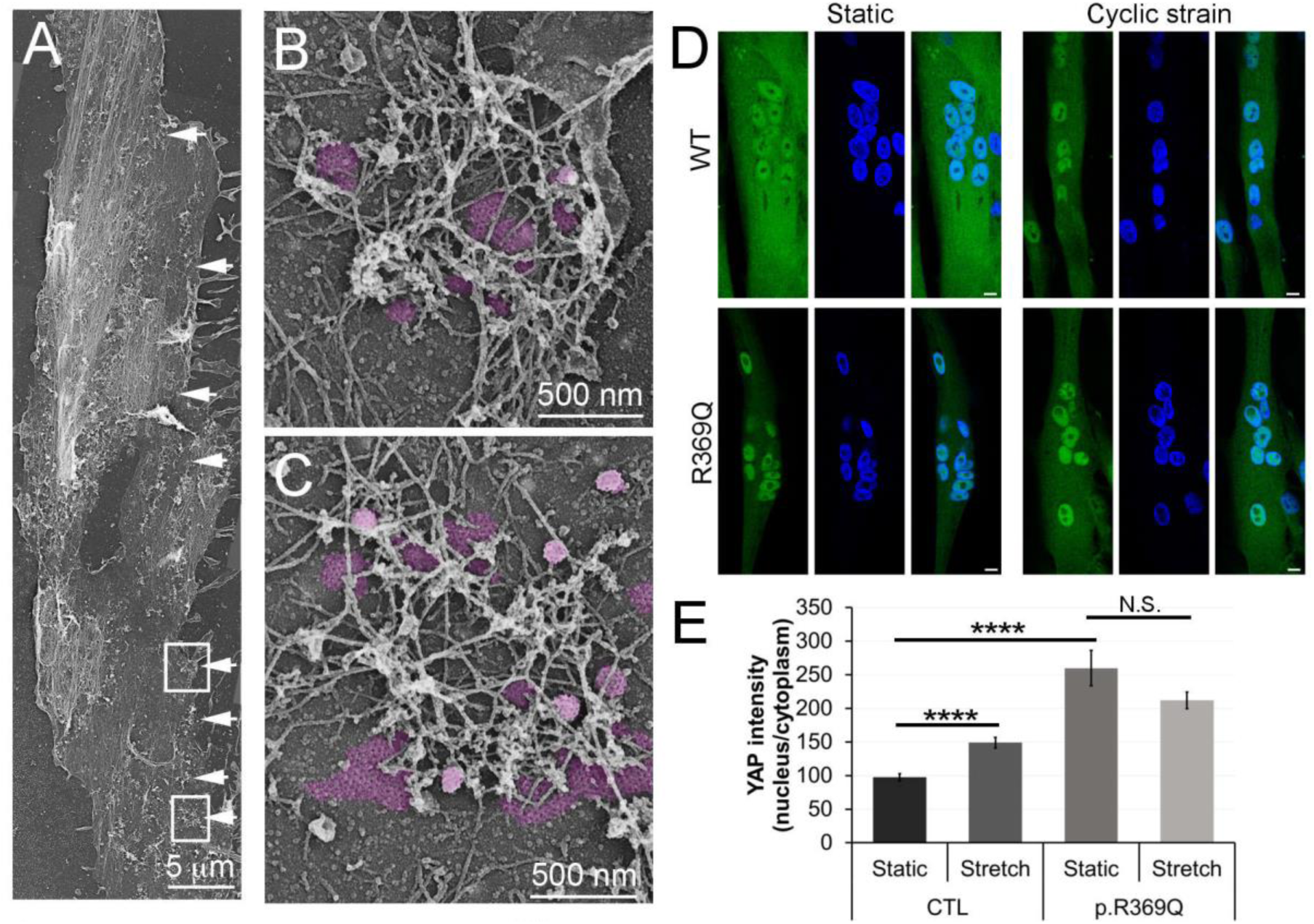
Human myotubes display extensive clathrin-coated plaques and CNM patient myotubes reproduce YAP signaling defects. **(A)** Survey view of the cytoplasmic surface of the PM from an unroofed human myotube. Arrows denote regularly spaced clathrin lattices. **(B) (C)** High magnification views of the boxed regions in (A). Clathrin lattices are pseudocolored in purple. **(D)** Immunofluorescent staining of YAP (green) and nuclei (blue) in control and p.R369Q myotubes either stretched (cyclic strain) or left static. **(E)** YAP fluorescence in control and p.R369Q myotubes either stretched (cyclic strain) or left static. (*n* = 35–60 myotubes). (Data presented as mean ± SEM; ****P < 0.0001).

**Supplementary Fig. 5.**
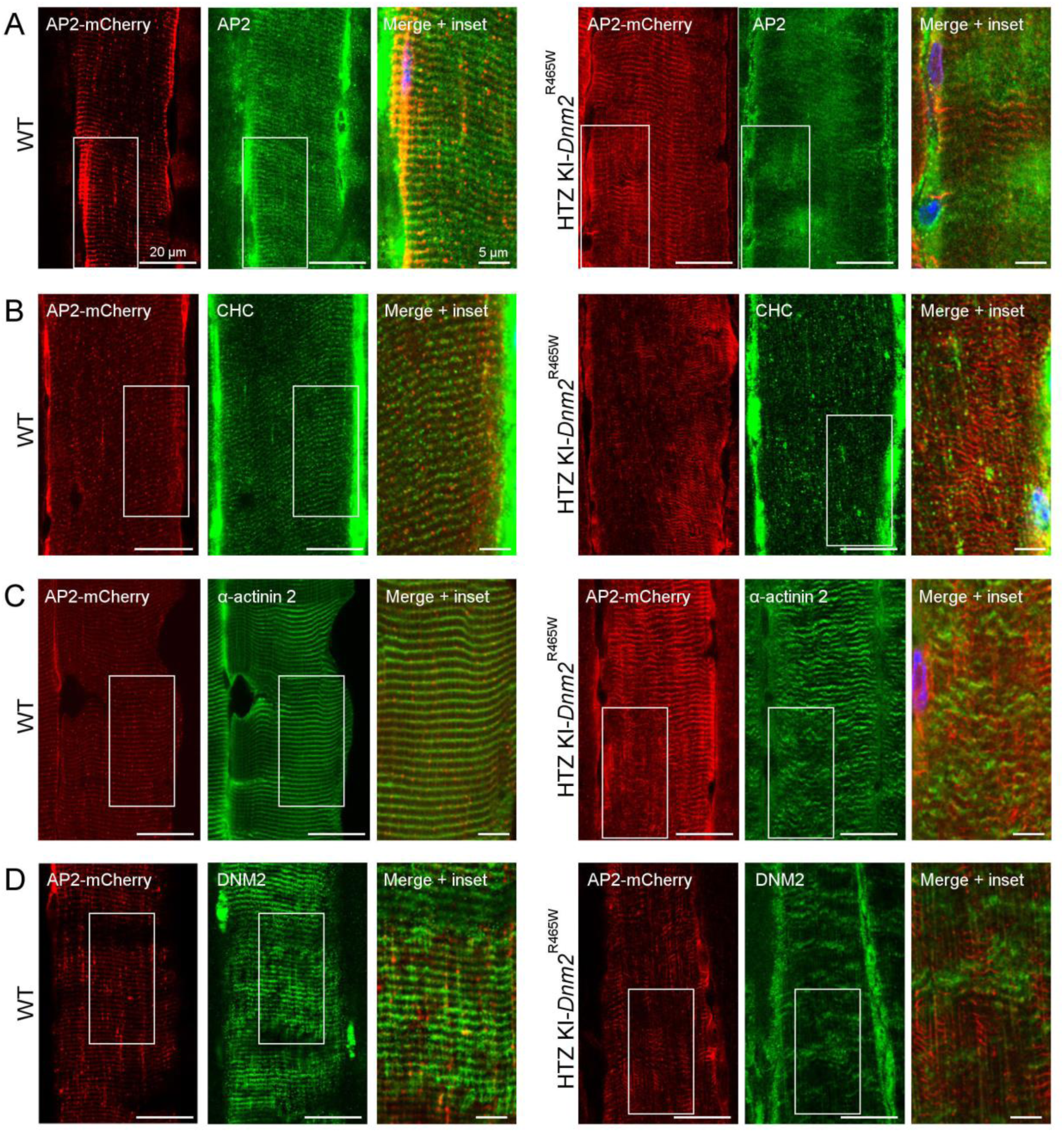
AP2-mCherry localization at skeletal muscle costameres. **(A-D)** AP2 mCherry distribution between WT (left) and HTZ KI-Dnm2^R465W^ (right) mice on fixed longitudinal muscle sections labelled with antibodies against endogenous AP2 (A), CHC (B), α-actinin 2 (C) or DNM2 (D) respectively.

**Supplementary Fig. 6.**
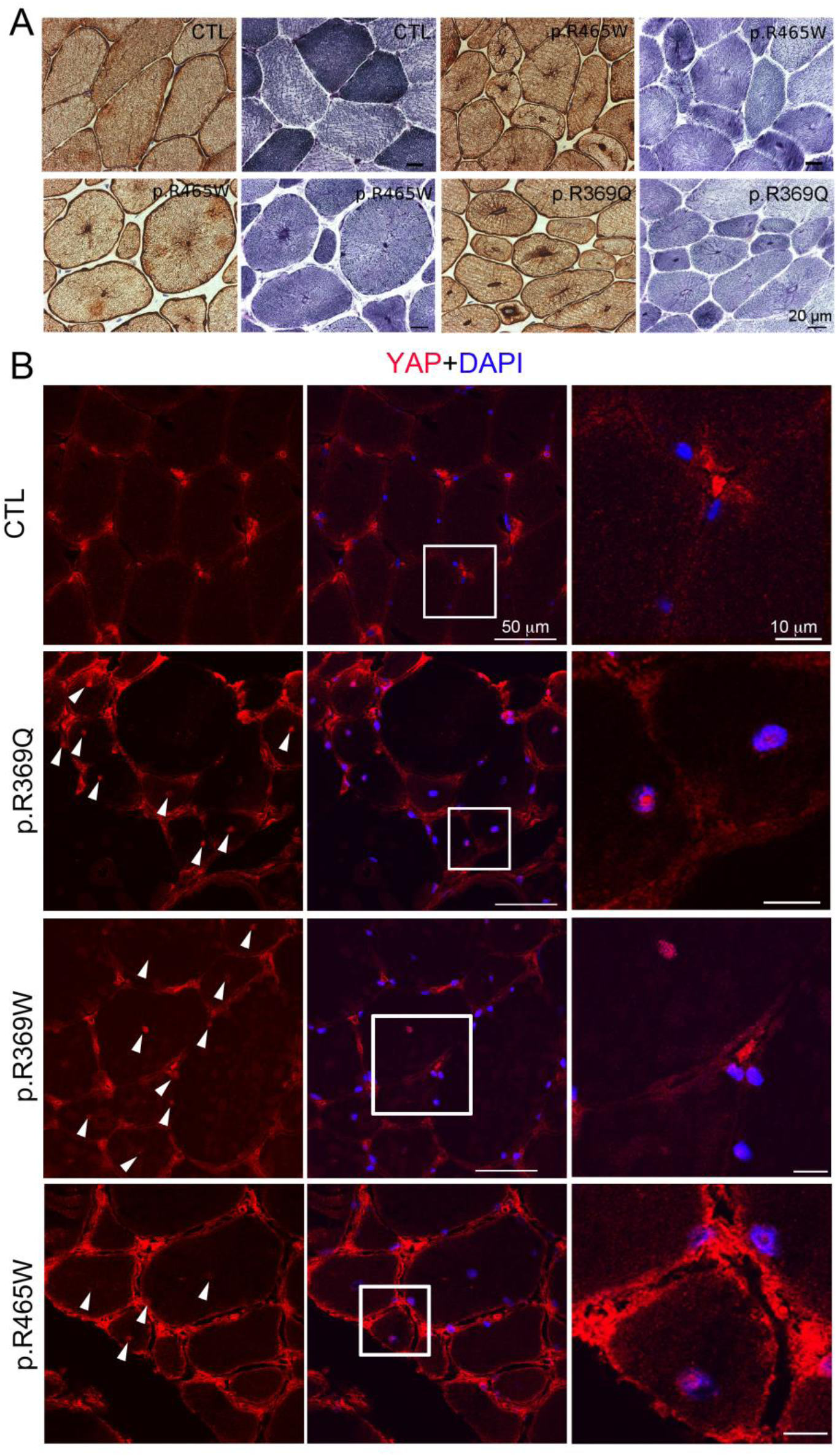
Desmin radial distribution and YAP-positive myonuclei in CNM patients. **(A)** Biopsy serial sections from one control subject (E-F) and three CNM patients (p.R369Q or p.R465W mutation). Immunohistochemical labeling against desmin (E, G, I, K) or NADH-TR reaction (F, H, J, L). Muscle sections reveal a strong radial sarcoplasmic strand desmin labeling. **(B)** YAP distribution in healthy skeletal muscle (CTL) and CNM patient (p.R369Q, p.369W, p.R465W mutation) skeletal muscle biopsies on transverse muscle sections labelled with antibodies against endogenous YAP (left panel) or merged with dapi.(middle panel). The right panel shows an inset of the boxed region in the merge.

